# Pharmacolibrary: Interoperable executable models for PK, PD and PGx derived from published evidence

**DOI:** 10.64898/2026.11.16.766960

**Authors:** Tomáš Kulhánek, Simon Denil, Burakcan Izmirli, Gökhan Ertaylan

## Abstract

Published pharmacological knowledge is distributed across heterogeneous scientific literature, databases, and computational models, making systematic reuse, comparison, and reproducibility difficult. We present Pharmacolibrary v26.11, a source-traceable resource that connects pharmacokinetic (PK), pharmacodynamic (PD), and pharmacogenomic (PGx) knowledge with executable models, simulations, and interactive visualizations. A multi-agent workflow combines LLM-assisted literature retrieval and extraction with deterministic validation, ontology mapping, provenance tracking, automated model generation, and rule-based review. The system transforms published evidence and extracted parameters into executable models across Modelica, FMI, SBML, MATLAB, and CellML and provides browser-based simulations for research and education. Preliminary evaluation indicates that decomposing literature extraction into multiple stages improves parameter recovery compared with single-pass extraction and that locally deployed open-weight models can provide performance that is at least comparable with frontier commercial LLMs. Pharmacolibrary provides a foundation for scalable, auditable curation of pharmacological knowledge and for connecting published evidence to reusable computational models across research, simulation, and education.

## Introduction

Computational and quantitative pharmacology integrates mathematical modeling, simulation, and data analysis with biological knowledge to characterize drug disposition, drug effect, and inter-individual variability. Its methods span educational compartmental models, population pharmacokinetic and pharmacodynamic (PK/PD) analyses, physiologically based pharmacokinetic (PBPK) models, and pharmacogenomic (PGx) investigations. Model-informed drug development and model-informed precision dosing have made such models central to both regulatory decision-making and individualized pharmacotherapy.(1,2)

Most of this modeling knowledge, however, remains poorly accessible, distributed across the text, tables, and figures of primary publications. Parameter estimates are reported in heterogeneous units and parameterizations, structural model assumptions are described in prose, and executable model code is rarely deposited in public repositories. Consequently, reusing a published model typically requires manual re-implementation, an error-prone step that limits reproducibility and prevents systematic comparison across drugs.

Several initiatives have addressed parts of this problem. The DDMoRe project and foundation (3,4) developed a tool-agnostic model markup language (PharmML) and a repository of curated PK/PD models; the project ended in 2016 and the repository is no longer maintained or publicly accessible (as of 2026), illustrating that long-term sustainability is as much a challenge as initial curation.

More recently, nlmixr2lib (5) has collected re-implementations of over 2 thousand published population PK/PD models covering over 1000 drugs distributed as model code for the open-source estimation package nlmixr2 (6). It is primarily intended to support model estimation by pharmacometricians working in R. In systems biology and physiology, BioModels (7) and the Physiome Model Repository (8) host curated models in the standard exchange formats SBML and CellML, respectively; clinical pharmacokinetics, however, is not their primary focus.

The Open Systems Pharmacology community maintains an open library of curated PBPK models together with software for model building, parameterization, and simulation (9). On the data side, PK-DB (10) provides human– and machine-readable access to curated pharmacokinetic data extracted from the literature and remains actively maintained. DrugBank (11) and T3DB (12) demonstrate that comprehensive drug and toxin metadata resources can be sustained over decades. ClinPGx, the successor of PharmGKB (13,14), curates drug–gene–phenotype associations and dosing guidelines. Finally, automated workflows for extracting pharmacokinetic parameters from the literature have begun to appear (15), suggesting a route past the manual-curation bottleneck.

Executable pharmacological models also have an important role in education. Pharmacokinetics and pharmacodynamics are challenging subjects because learners must connect physiological processes, mathematical parameters, drug dose, concentration–time profiles, and pharmacological response. Simulation and visualization are increasingly used to support PK education (16), including recent open-source tools such as PK-SIMple that aim to make pharmacokinetic simulation more accessible for teaching (17). These applications illustrate that executable models can serve not only as reproducible computational artifacts for researchers, but also as interactive learning objects that can be explored and modified to connect drug behavior with underlying physiology. For this purpose, models benefit from being accessible across different software environments and usable without specialized pharmacometric expertise.

What is still missing is a resource that couples curated PK, PD, and PGx parameters, each traceable to its source, with executable models covering a broad range of drugs, while making those models usable beyond the pharmacometric community. Systems biologists, biomedical engineers, and physiologists developing integrative models of human physiology may work outside the R, NONMEM, nlmixr2 or Open System Pharmacology ecosystems and require portable drug models that can be coupled to their own physiological models. Educators may similarly benefit from models that can be readily explored and simulated without extensive software installation or programming. Such a resource must also support incremental and sustainable curation rather than rely on one-off manual implementation.

For industry, such a resource would reduce the repeated effort of reviewing and rebuilding drug models from the literature in each project: a library of source-traceable and executable models can serve as a starting point for model-informed drug development, which is already used across the drug lifecycle and in regulatory decisions (18). Traceability of every parameter to its source also supports model credibility assessment (19), and an open, standards-based format makes models easier to share with partners and reuse in different tools, as the OSP community has shown for PBPK (9).

As a first step toward such coverage, Pharmacolibrary v25.08 (20) provided reusable Modelica components for PK, PD, and PGx modeling together with automatically generated PK models of almost all drugs organized according to the Anatomical Therapeutic Chemical (ATC) classification. Modelica is an open, equation-based, object-oriented language for modeling complex dynamic systems, in which models are composed from reusable components connected through standardized interfaces and can be simulated in both open-source and commercial tools (21). The Pharmacolibrary follows CalVer (Ubuntu-style) calendar versioning (vYY.MM), where the version identifies the year and month of release rather than a sequential version number as multiple improvements and especially generated models of drugs are anticipated to be updated regularly in future months and years. The library formalized established pharmacological terms, compartments, and processes as reusable modeling components. Some of its models were also deployed as interactive web-based simulations using the Bodylight.js toolchain (22,23), including examples in which pharmacological effects were coupled to physiological models to demonstrate drug-induced changes in a modeled physiological system (20).

The Pharmacolibrary version v25.08 has four important limitations. First, the LLM was used to extract knowledge from some of the relevant journal articles and from the general knowledge of the LLM itself to produce PK models, but it had not been systematically validated against the corresponding published results, limiting confidence in their reproducibility and reuse. Hallucination and reliability is one of the big concerns when using AI and LLM for this type of tasks. Second, the v25.08 models primarily represents population-level or reference individuals and do not explicitly capture inter-individual variability through covariates reported in scientific publication and clinical trials. In applications related to individualized pharmacotherapy, drug exposure and response may depend on characteristics such as age, body weight, sex, ancestry, hepatic and renal function, cardiac output, and genotype. Explicit representation of such covariate relationships allows model predictions to be conditioned on individual characteristics and can reduce unexplained variability. Third, the v25.08 library is limited to the Modelica modeling environment, restricting direct reuse by communities working with other modeling and simulation technologies. Fourth, only population pharmacokinetics is systematically presented among all drugs that are in the ATC classification system.

Here, we present Pharmacolibrary v26.11, a substantially extended version that integrates pharmacokinetic, pharmacodynamic, and pharmacogenomic knowledge with executable models and simulation, while providing interactive access to the evidence underlying the structured knowledge. The new release extends the original Modelica library v25.08 (20) into four coupled components: (i) a machine-readable and human-explorable knowledge base of PK, PD, and PGx parameters, with each record linked to its source publication; (ii) a repository of corresponding executable models in multiple modeling and exchange technologies, including Modelica, FMI, SBML, MATLAB, and CellML, accompanied by scripts for simulation and replication; (iii) a collection of ready-to-run web-based simulations of selected models; and (iv) knowledge-base database compressed and presented as an interactive chat example application. Pharmacology– and physiology-relevant parameters, including explicit covariate relationships, are extracted from the primary literature using an automated LLM-based pipeline. Because automated extraction cannot be assumed to be error-free, records retain source provenance and curation status, while the workflow identifies cases requiring further human review based on extraction or model-validation results.

Because the long-term maintainability is of concern, all the Pharmacolibrary is shipped as a static web page with all computation and querying executed in-browser, thus not relying on server side capabilities. The remainder of this article describes the knowledge-base architecture, literature-extraction and quality-control workflow, model generation and validation, supported modeling and exchange technologies, and evaluation.

## Methods

We implemented a multi-agent architecture to populate and maintain the knowledge base and its associated computational artifacts. The high-level software architecture of the agents and supporting system is shown in Figure 1. The workflow comprises three main agents—Scholar, Engineer, and Reviewer—which operate on a shared repository and iteratively refine the extracted data, executable models, and simulation results.

**Figure 1:**
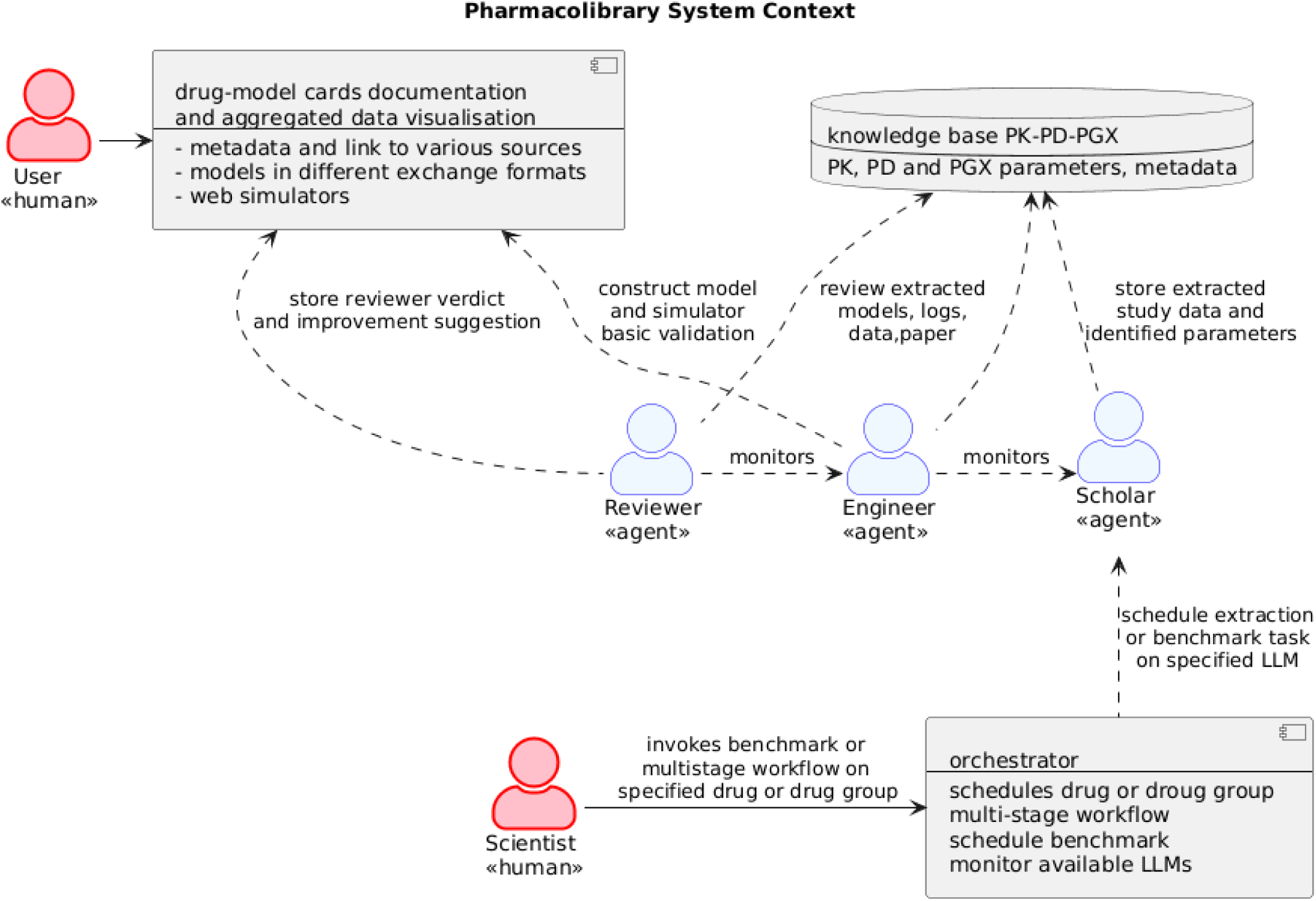
Overview of the multi-agent architecture used to populate and maintain the Pharmacolibrary knowledge base. The Scholar agent retrieves and extracts PK, PD, and PGx information from available literature and supporting artifacts; the Engineer agent generates executable models and simulations; and the Reviewer agent evaluates the extracted data and model behavior against the source evidence. Extracted data, model metadata, provenance, and simulation results are stored in a shared repository, enabling iterative refinement and identification of artifacts requiring human review.

The Scholar agent has access to tools that can directly query for publication, download abstracts or full papers of the relevant scientific publication. It focuses on a defined topic, such as the pharmacokinetics of a specific drug, and searches online literature and data resources for relevant information. Based on the search results, it retrieves available source artifacts, including open-access publications in PDF or Journal Article Tag Suite (JATS) format (24), as well as supplementary materials such as source code, tables, and additional text. Retrieved artifacts are processed by an extraction pipeline that filters irrelevant content and identifies, transcribes, normalizes, and validates information relevant to the requested topic, such as PK, PD, or PGx parameters. Each extracted record is linked to its source artifact and location, providing provenance for subsequent processing. When parameters required for model construction are not explicitly reported in a given source, complementary information may be obtained from other artifacts associated with the same study or drug. Such gap-filled values are explicitly marked as inferred or supplemented rather than treated as directly reported measurements. For example, an executable oral PK model may require bioavailability, absorption rate, or lag time even when one or more of these parameters are not reported in the primary publication.

The Engineer agent has access to a tool that can operate on the parameter sets and structural information extracted by the Scholar agent and generates executable models in the supported modeling and exchange technologies. Generated models undergo automated execution checks to verify that they can be compiled or instantiated and simulated successfully, produce the expected model outputs such as concentration–time or effect–time profiles, and do not exhibit obvious numerical or physiological inconsistencies. These checks provide an initial level of model quality control but do not establish agreement with the published study.

The Reviewer agent evaluates the generated models against the source publications. Where a publication reports simulation results graphically or numerically, reference data are extracted from tables or digitized from figures and compared with the corresponding model simulations within predefined tolerances. The reviewer records discrepancies in model structure, parameterization, or simulated behavior and produces a review report that can be used to trigger further refinement by the Scholar or Engineer agents or to flag the artifact for human review.

### Scholar agent — knowledge-base scaffolding and literature extraction

Scholar agent has access to tools that can search and download appropriate data available via the Internet either by himself or indicate to be prepared by the human user. The knowledge base is scaffolded from information retrieved from established drug and pharmacology resources, including DrugBank(11), T3DB(12). The Anatomical Therapeutic Chemical (ATC) classification system (25) is used to associate drugs with ATC codes and names. The clinical pharmacogenomics resource ClinPGx (26) is used to identify drug–gene–phenotype relationships and relevant pharmacogenomic guidelines. The initial drug list is further expanded using metadata available through the European Medicines Agency (EMA) portal (27).

These sources provide the initial structure of the knowledge base, which is organized by drug and, where applicable, registered drug combinations. Additional PK, PD, and PGx information is subsequently extracted from the scientific literature using the multi-agent workflow described in Figure 1. The Scholar agent then performs literature retrieval and extraction, while the Engineer and Reviewer agents use the resulting records to generate and assess executable models.

The Scholar agent implements a deterministic sequence of retrieval and extraction steps, with LLM-assisted processing used for relevance assessment, localization, transcription, interpretation, and validation of the extracted information.

First, the agent retrieves publications relevant to the target drug and topic, such as population pharmacokinetics, pharmacodynamics, or pharmacogenomics. For each identified publication, bibliographic metadata, including title, authors, abstract, and persistent identifiers, are collected. For openly accessible publications, the agent attempts to retrieve the full text and associated supplementary materials, including source code, tables, and additional text.

The ‘retrievè step uses the PubMed and Europe PMC APIs to search and identify relevant publications. For pharmacokinetic studies, queries combine the drug name or known synonyms with terms such as “population pharmacokinetics”; analogous topic-specific queries are used for PD and PGx. Query breadth can be adapted when an initial search retrieves too few relevant publications or when subsequent extraction steps fail to identify the required information. Semantic Scholar (28), Unpaywall(29), Europe PMC(30), and OpenAlex(31)are used to identify and resolve openly accessible full-text sources. Where publishers provide direct access to open-access content, the article is retrieved from the publisher. Publications for which automated retrieval is not possible are flagged for potential manual acquisition, for example through institutional subscriptions, and can subsequently be processed by the Scholar agent. Whenever available, structured article representations in Journal Article Tag Suite (JATS) format (24) are preferred because they preserve the logical structure of the publication. If JATS is unavailable, full-text content is extracted from PDF documents using GROBID (32) and PDFPlumber as fallback service to read PDF documents (33). Figures are processed separately using the vision capabilities of the LLM to identify and transcribe information that is not available in the extracted text, including graphical simulation results and parameter values. The development version of this agent implemented a “single-pass” extraction workflow of the Scholar agent; it uses a single prompt containing the abstract or full text retrieved in order to extract PK, PD, and PGx parameters. This approach produced unsatisfactory extraction quality and systematic errors that were difficult to address through prompt engineering alone – as reported later in Table 3. Therefore a second, “multi-stage” pipeline workflow version of the scholar agent decomposes the extraction process into separate stages. The purpose of this decomposition is not only to extract parameter names and values, but also to establish explicit criteria for assessing parameter quality, provenance, and interpretation. Each stage uses a text-capable and, where required, vision-capable LLM to perform a specific task. Failure of an individual stage terminates the extraction workflow for that artifact and records the corresponding failure reason.
The ‘screen’ step assesses the relevance of a publication by estimating whether its content contains extractable population PK, PD, or PGx information for the target drug. Publications are accepted when they contain model parameters or other structured information relevant to the target task and rejected when the target drug is only mentioned, for example as a co-administered comparator, while the reported quantitative data refer to another drug.
The ‘locatè step identifies the sections, tables, figures, or supplementary materials most likely to contain the relevant model parameters and distinguishes parameter tables from unrelated content such as demographic or baseline-characteristic tables. It also identifies the locations of standard NONMEM parameters and their definitions, which are frequently reported in supplementary materials.
The ‘transcribè step transfers the content of each located region into a structured typed record without assigning additional semantic interpretation. For example, a table entry labelled “V” with a value of “183” and unit “mL” is transcribed as reported. Information extracted from figures is similarly retained together with its source location.
The ‘interpret’ step assigns semantics to the transcribed records. Parameter labels are mapped to canonical concepts using an extended PK ontology based on the previously described PK ontology (15). The step identifies parameter types and covariate relationships, resolves references to NONMEM THETA indices, and distinguishes apparent parameters from structural parameters and compound-specific identities.
The ‘validatè step applies rule-based consistency checks to the extracted records. These checks include unit consistency, arithmetic relationships between reported quantities, detection of missing structural parameters, and plausibility checks against predefined constraints. For example, where volume of distribution, clearance, and half-life are available, the reported half-life is compared with the value implied by the corresponding clearance and volume parameters.
The ‘crosscheck’ step repeats the LLM dependent stages (screen, locate, transcribe, interpret and validate) using a genuinely different LLM model and compares the resulting structured records with the primary extraction. Agreement and disagreement are determined by deterministic comparison rules, and disagreements are recorded for subsequent review.

For each drug, the Scholar agent produces a set of parameter records containing the original label reported in the publication, its canonical semantic class, value, unit, normalized value and unit, uncertainty information where available, and a provenance locator identifying the exact source region. An example is shown in Table 1.

**Table 1:**
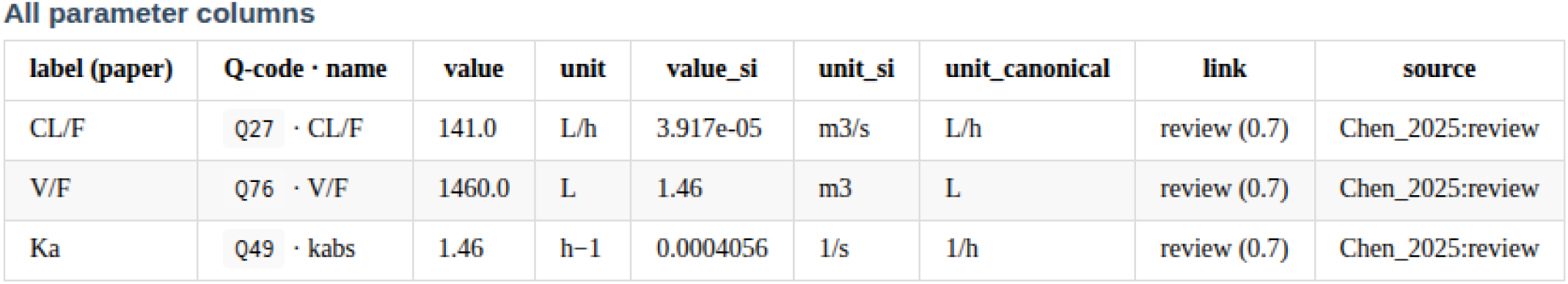
Example of PK parameter records extracted by the Scholar agent from a publication reporting the pharmacokinetics of metformin (34). Each record retains the original parameter label, canonical parameter concept, reported and normalized values, unit, and source locator.

The extraction, validation, and cross-check status of each record is retained as structured metadata and presented to users through status badges and brief explanations when discrepancies or failures are detected.

### Engineer agent – executable model generation and simulation

The Engineer agent consumes ontology-linked PK, PD, and PGx records selected by the Scholar agent for model reconstruction. It deterministically translates the extracted structural parameters, covariate relationships, and model topology into executable implementations using multiple modeling and simulation technologies. These include Modelica as the canonical modeling language, MATLAB for executable implementations and simulation scripts, FMI for model exchange and co-simulation, and SBML and CellML for interoperable model exchange format.

Generated models are simulated using OpenModelica tool (35) to perform basic execution and plausibility checks and to produce concentration–time or effect–time profiles. From the simulated profiles, commonly reported PK measures such as C_max, T_max, AUC, and, where applicable, terminal half-life are calculated. The Engineer agent also generates the input configuration required by the web-based simulator. When a parameter required by a model template is not available in the source data, an explicit base-model default may be used. Such substitutions, together with structural assumptions and interpretation flags, are recorded in a model provenance card and deviation report. The parameter estimates supplied by the Scholar agent are otherwise preserved without modification.

The canonical modeling components of Pharmacolibrary v26.11 implemented in Modelica are shown in Figure 2A. Detailed definitions of the components, selected connectors, and quantities exchanged between components were previously described for Pharmacolibrary v25.08 (20) and remain the basis of the current implementation. These components can be combined to construct executable pharmacological models using the corresponding governing equations.

**Figure 2:**
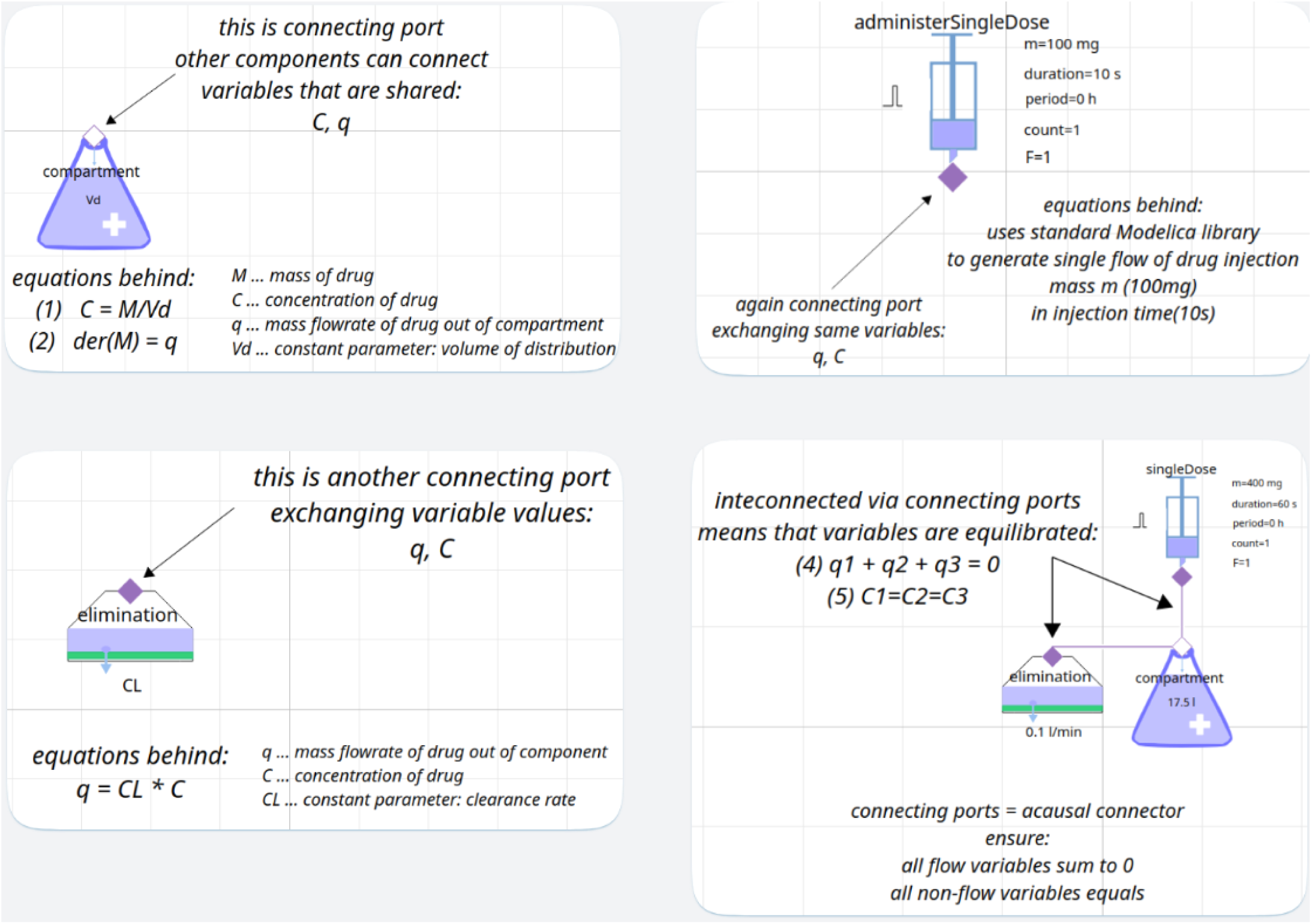
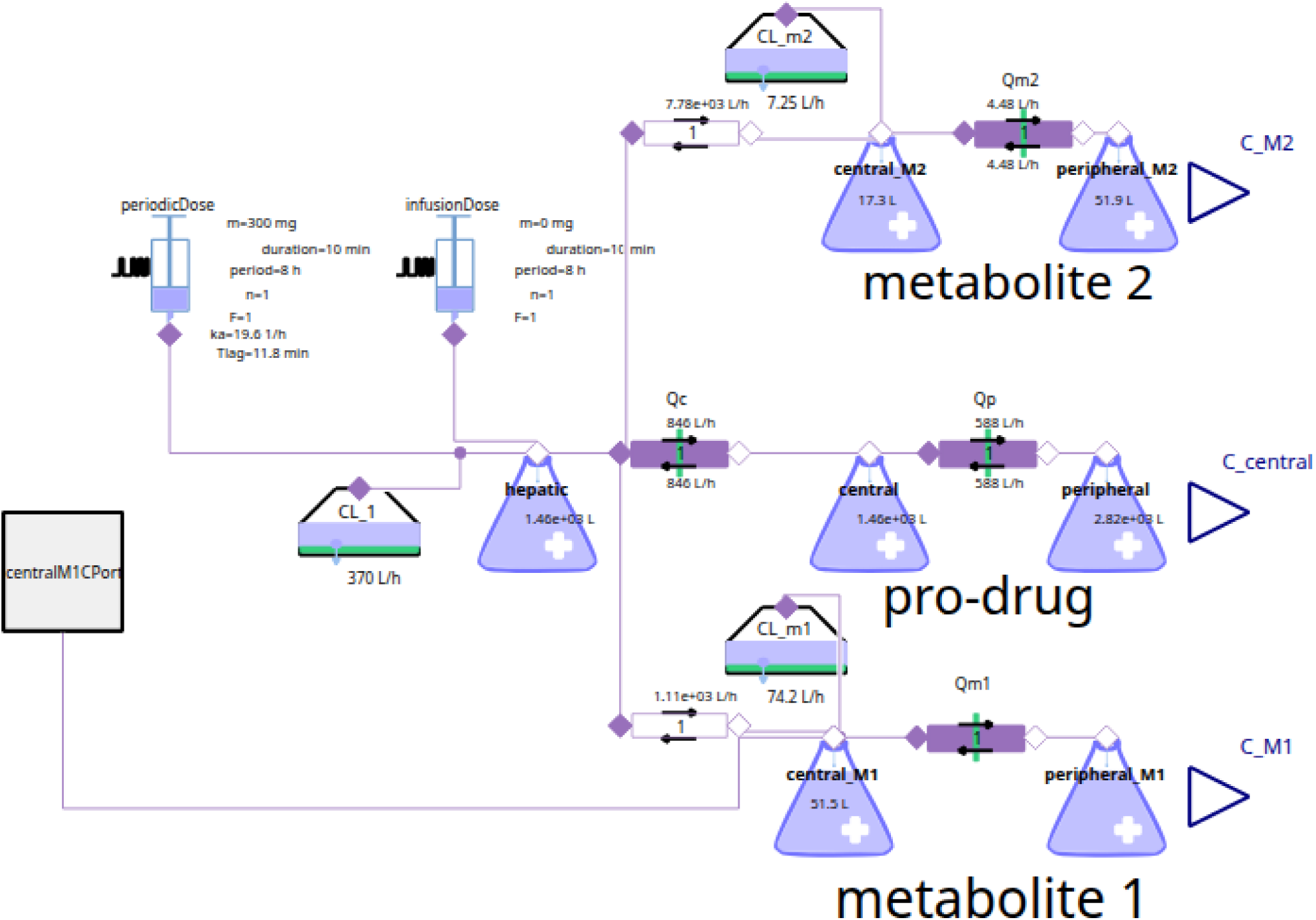
(A) Canonical Pharmacolibrary v26.11 modeling components implemented in Modelica, including their governing equations. Components can be connected to construct executable pharmacological models. Detailed component definitions, connectors, and quantities are described in (20).(B) Generated Modelica model of clopidogrel pharmacokinetics and its metabolites, rendered by OpenModelica (35), with parameters extracted from Jung et al. (36). The model includes the H4 active metabolite (metabolite 1) and the carboxylic acid metabolite (metabolite 2).

Figure 2B shows an example generated model of the pharmacokinetics of clopidogrel and its metabolites, rendered by OpenModelica(35), using parameters extracted from Jung et al. (36). The model represents the H4 active metabolite and the carboxylic acid metabolite.

### Reviewer agent — rule-based verification of extracted and replicated models

The Reviewer agent provides an independent downstream assessment of the outputs produced by the Scholar and Engineer agents. It evaluates the Scholar’s interpreted records and source transcriptions together with the Engineer’s model, simulation results, and deviation report. Final pass/fail decisions are deterministic and rule-based. LLMs are used only for auxiliary tasks such as checking whether deviations are adequately documented and generating human-readable review notes, and cannot alter the decision.

For each paper–population record, the Reviewer evaluates four aspects: (T0) analyte identity, ensuring that the modeled compound corresponds to the reported analyte; (T1) replication, comparing simulated PK measures such as C_max, T_max, and terminal half-life with reported values using predefined agreement criteria; (T2) covariate representation, checking whether reported covariate effects are represented in the model or tested scenarios; and (T3) construction consistency, covering model structure, output variables, parameter coverage, apparent parameters such as CL/F, unit and rate-constant conversions, and documentation of defaults or structural assumptions.

The individual checks are summarized into measures of provenance completeness, internal consistency, extraction agreement, replication fidelity, and construction fidelity. All records are cross-checked using a genuinely different second LLM, and the reviewer assesses extraction agreement by comparing the primary and cross-check outputs. The resulting assessment is assigned a rule-based status such as *curated (human checked), reproduces paper, reproduces with caveats,simulated, not simulated, needs review, rejected*. Promotion to a curated record remains a human decision. An example of such a Reviewer’s verdict is in Figure 3 showing caveats and explanation of it.

**Figure 3:**
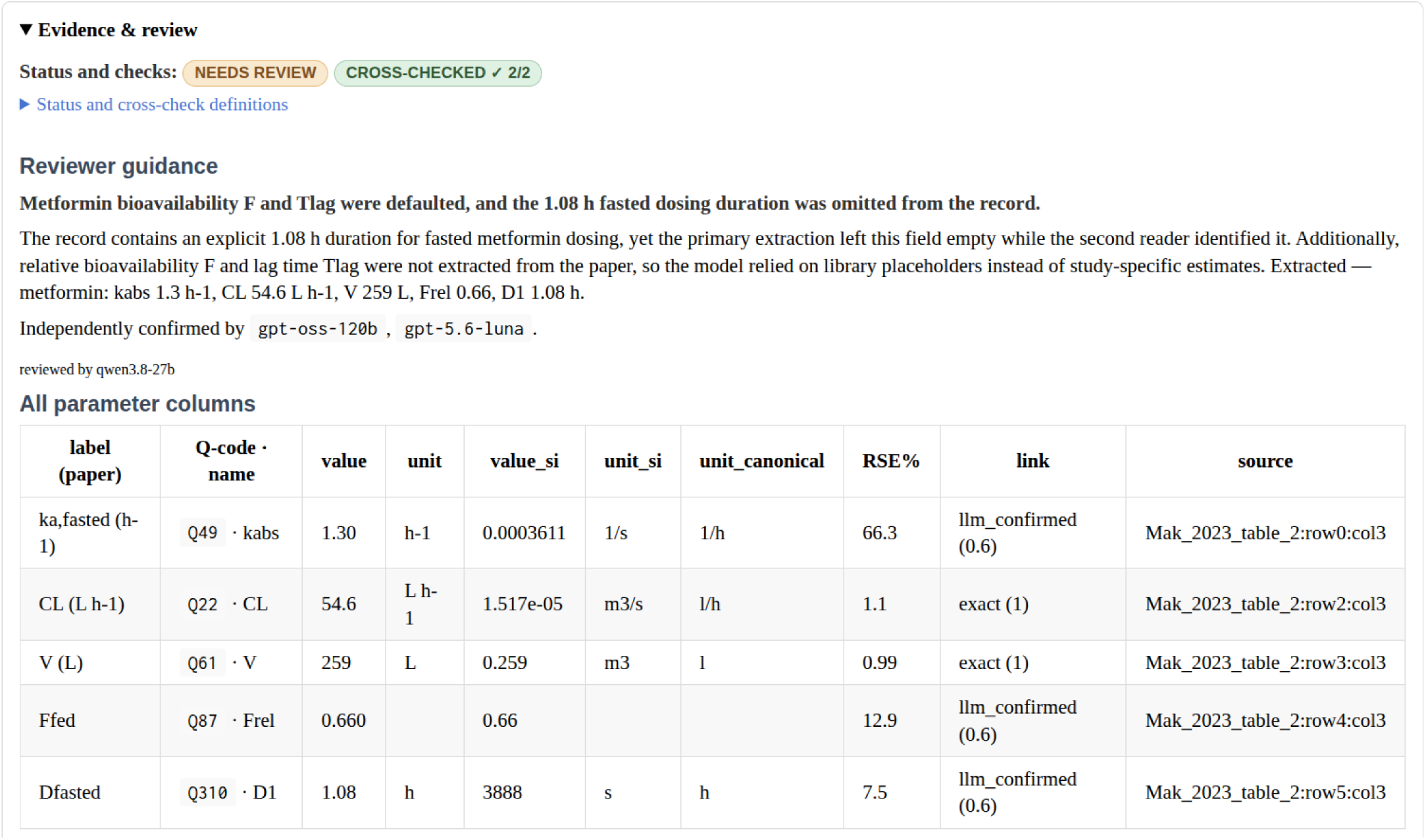
Reviewer output for the metformin population-PK models of Mak et al. (37). Both model implementations were accepted with caveats: the structural model was reproduced, whereas the reported fasted dosing effects were not modeled and simulated. The figure illustrates the rule-based verdict, supporting evidence, and independent extraction-agreement assessment.

**Figure 4:**
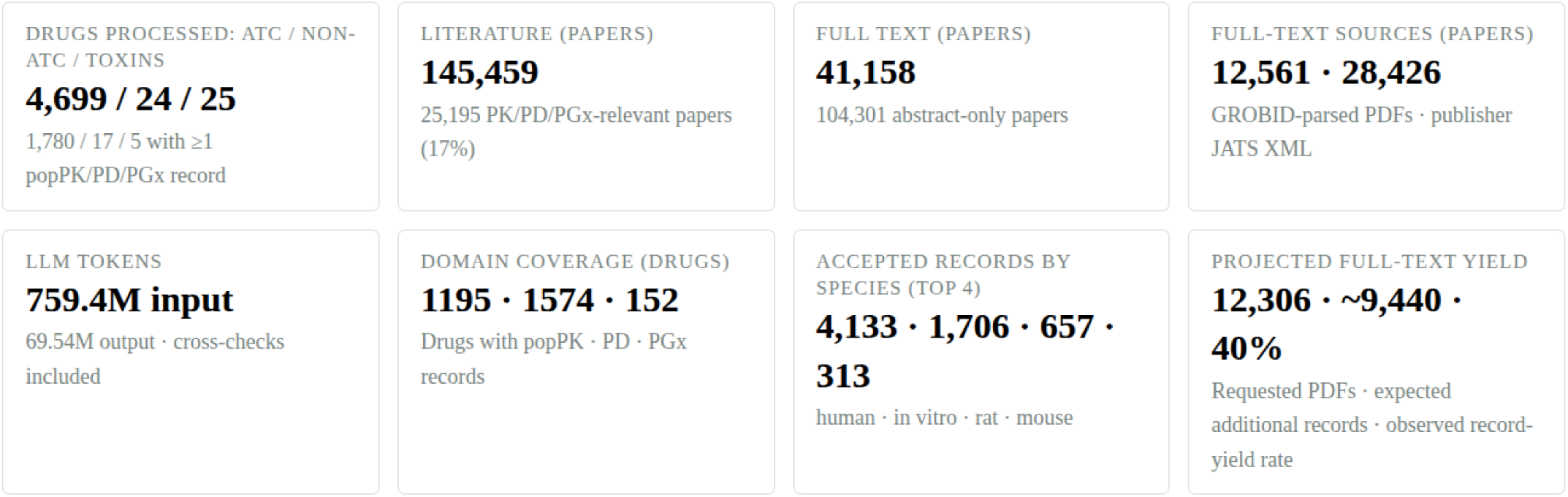
Statistics of extracted information of drugs and compounds and about their popPK,PD,PGx records.

Each record receives a concise explanation of the verdict, with the relevant parameter values and source evidence. At the drug-group level, the Reviewer additionally summarizes verdict distributions, recurrent blocking reasons, extraction agreement, and recurring workflow issues. These group-level findings are used to identify improvements for the iterative development/evolution to the Scholar and Engineer workflows while remaining explicitly separate from the curation decision.

On top of the all agent’s tasks, orchestrator creates three SQLite databases: a query database with the extracted records (∼7 MB), a passage database of DrugBank, ClinPGx and abstract text for the chat’s model (∼26 MB), and an ADME database(∼7MB) containing tissues/site of action. Each file is built in a fixed order, compacted, and named by a hash of its content, so an unchanged knowledge base gives the identical file and the browser can cache it indefinitely. The passages are short, sentence-bounded chunks with a full-text index, and the passage database loads only when a reader switches to “Ask the data” page and selects to load a language model.

## Results

To date, 4,683 ATC-classified drug compounds, 24 non-ATC drug compounds, and 25 toxins have been processed using the multi-stage workflow. The workflow screened 134,781 publications, of which 37,279 were obtained as full texts and 97,502 were screened using abstracts only. Of these publications, 23,229 were judged relevant for the extraction of population PK, PD, or PGx study data. Approximately 10,114 article PDFs were parsed using GROBID (32) to obtain structured full-text information, while most full texts (27,041) were obtained in publisher XML (JATS) format. The multi-stage workflow consumed 652 million input tokens and generated almost 51 million output tokens.

Not all drugs had extractable population PK, PD, or PGx information in the retrieved literature. However, at least one population PK record was extracted for 947 drugs, at least one PD record for 1,690 drugs, and at least one PGx record for 150 drugs. Human studies accounted for the largest group of records (3,903), followed by in vitro studies (1,694), rat studies (623), and mouse studies (296).

The workflow also maintains a “wanted PDFs” list of publications whose full texts could not be obtained because of paywalls or other access barriers, but whose abstracts suggest a high probability of containing relevant PK, PD, or PGx data. This list currently contains 12,306 publications, which could potentially yield an additional 9,440 records. This estimates comes from ratio of 40%, which is number of extracted records from highly relevant scored abstract of open-access resources.

To date, eight distinguishable model templates have been selected to cover pharmacokinetics of majority reported drugs, including single– and multicompartment models and composite models following the parent–metabolite pattern. Thus for every specific PK model instance, the engineer agent chooses the most appropriate model template, inherits its structure and specifies only different parameter values. Additionally PK-driven or dose-exposure templates were selected to cover reported pharmacodynamics. The object-oriented structure of the templates allows to keep the number of structural model templates relatively low and every specific PK,PD model is instantiated with an individual parameter set and exported to a target modeling or simulation technology that was not necessarily used in the original publication. This enables reconstruction of the published model structure while preserving the extracted parameterization and documenting deviations introduced during model generation. The canonical Modelica components provide the reference representation from which the supported executable and exchange implementations in other formats are generated. An example of the Engineer agent output for an extracted metformin PK parameter set is shown in Figure 5.

**Figure 5:**
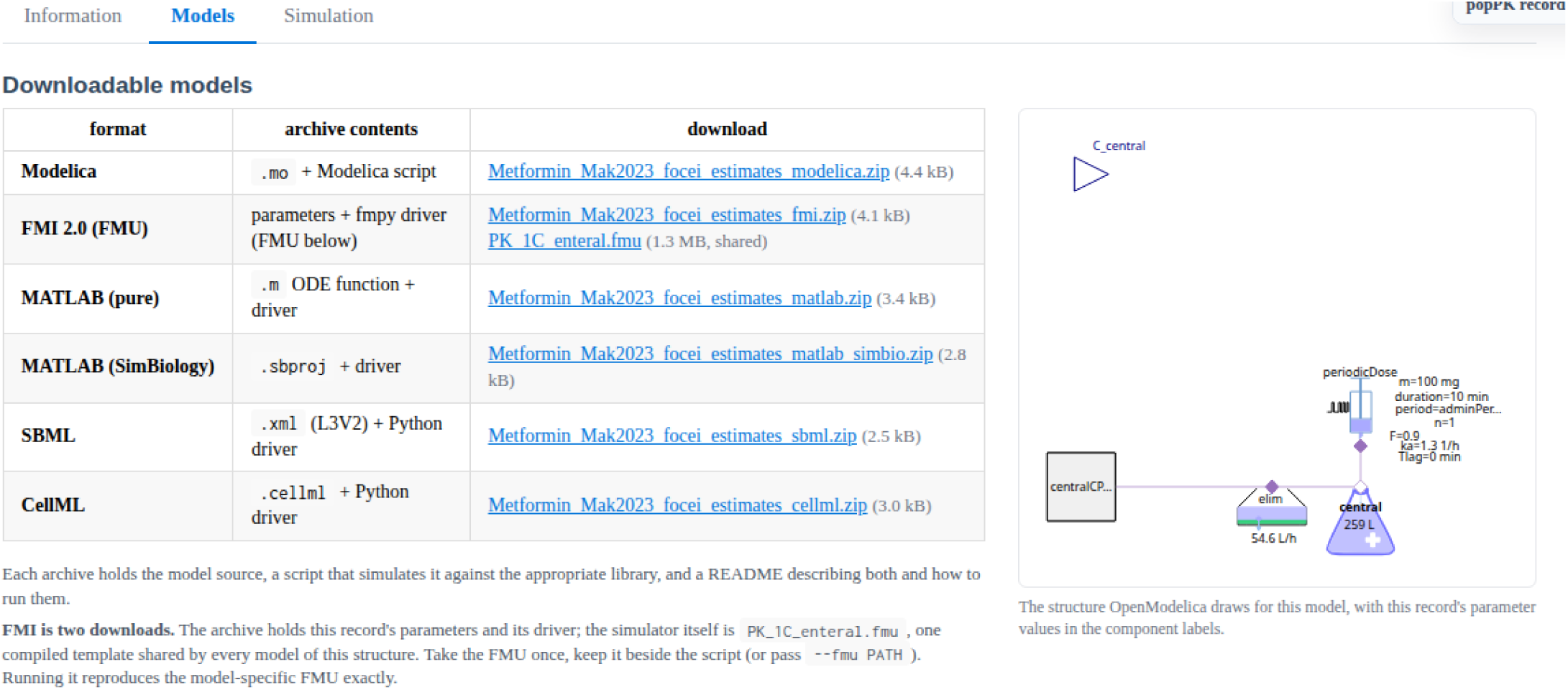
Examples of generated model implementations in various modeling and exchange formats for extracted metformin PK parameter set reported by Mak et al.(37)., including the corresponding one-compartment model diagram rendered by OpenModelica (35).

The Engineer agent generates a web-based simulator from the executable model. Using FMI-based model packaging and the Bodylight.js toolchain (23), the appropriate template model is compiled to WebAssembly and integrated into a client-side interface instantiated with drug/paper specific parameters. The resulting simulator executes the model locally in the web browser without requiring simulation on a remote server. Parameter values are initialized using the values extracted from the source publication, while selected parameters can be modified interactively and the simulation rerun to visualize the resulting changes in drug concentration curve. Figure 6 compares a web-based simulation of pharmacokinetics of metformin using the extracted parameters with an updated simulation generated immediately after the user modifies selected parameters. Figure 7 shows a web-based simulation of the exposure–response relationship for the antithrombotic agent clopidogrel using the extracted pharmacodynamic parameters. Figure 8 shows a web-based simulation of pharmacogenomic effects on the pharmacokinetics of clopidogrel and two of its metabolites.

**Figure 6:**
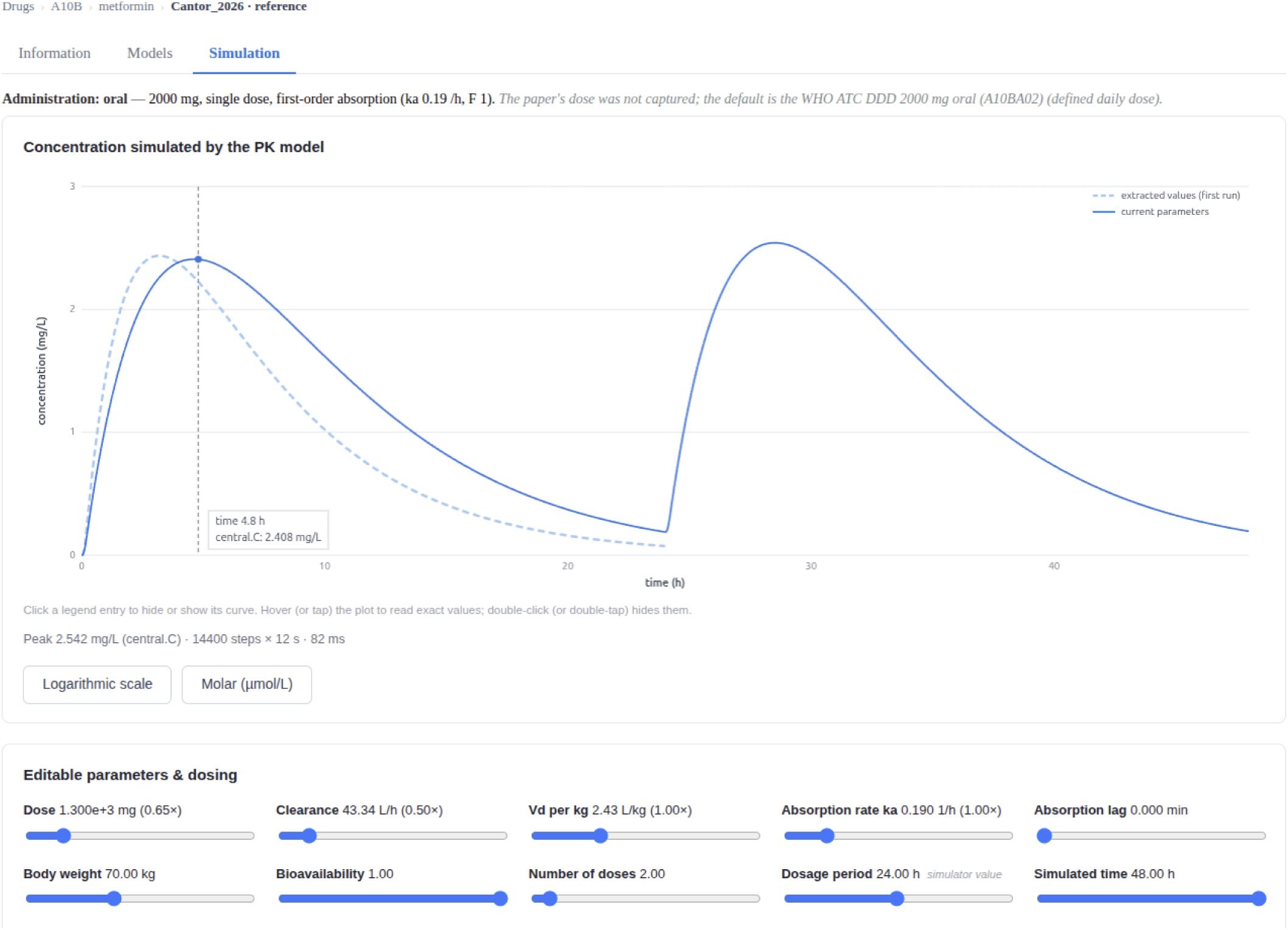
Example of a generated web-based simulator for the extracted PK model of metformin. The light blue dashed curve shows the default simulation with parameter values initialized from the published study; the dark blue curve shows the simulation after the user interactively reduced clearance to 50% of its original value, reduced the daily dose to 1300 mg and doubles the simulation time and adds 2 doses each administered every 24h.

**Figure 7:**
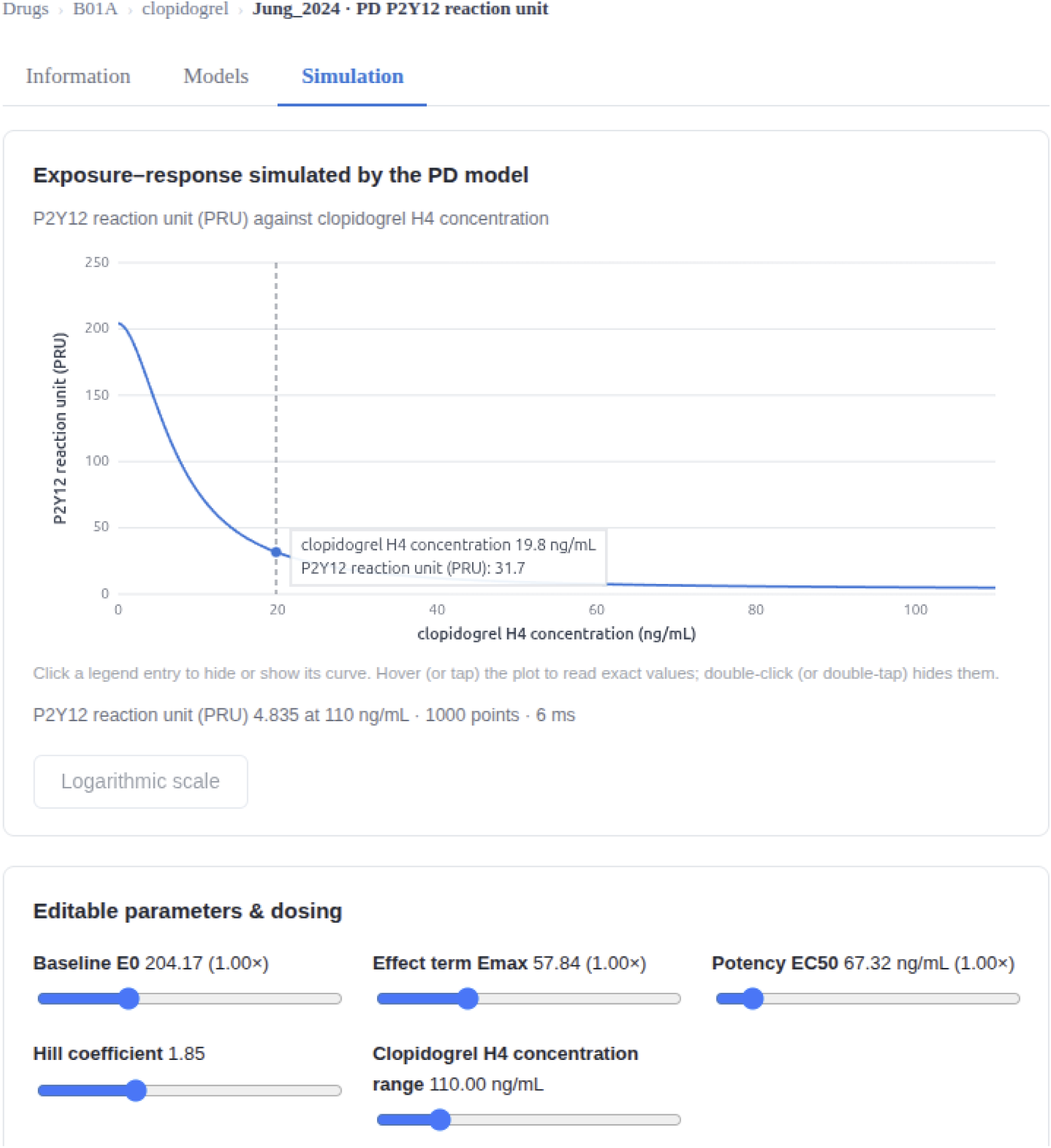
Example of a generated web-based simulation of an extracted PD model of clopidogrel. The exposure–response curve, based on the extracted Emax model parameters, shows the decline in P2Y12 reaction units (PRU) with increasing exposure to the drug.

**Figure 8:**
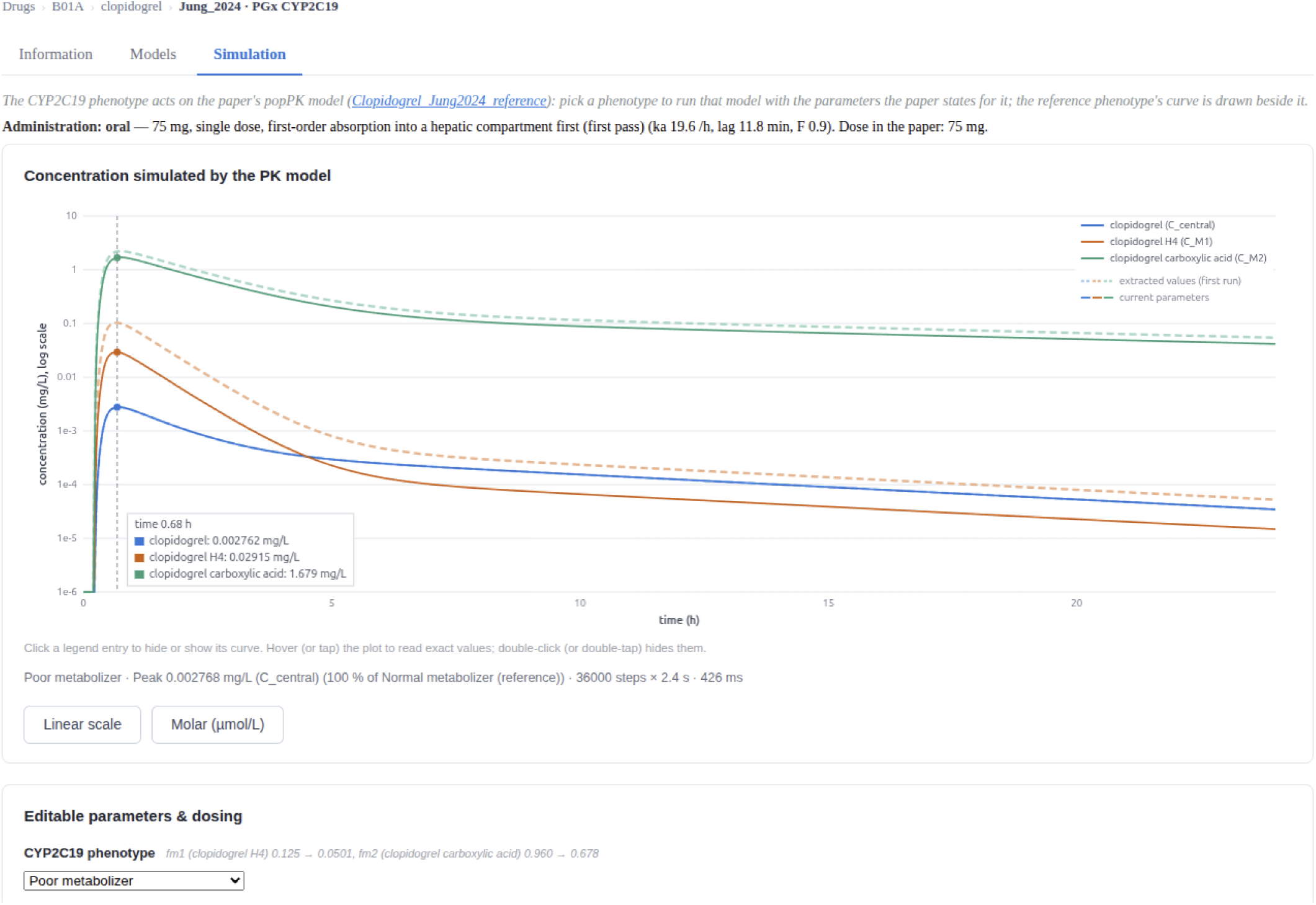
Example of a generated web-based simulator of a PK model with PGx influence extracted from a published study, showing the influence of CYP2C19 phenotype on the pharmacokinetics of clopidogrel and two of its metabolites. In a simulated patient whose pharmacogenomic profile indicates the CYP2C19 poor metabolizer phenotype, plasma concentrations of clopidogrel are similar to those in a normal metabolizer, whereas the active metabolite H4 and the carboxylic acid metabolite reach lower levels (due to poor metabolic effect of CYP2C19 enzyme) over the dosing interval, which is expected to reduce the antiplatelet effect in this patient. Curves for a normal metabolizer are shown as dashed lines for reference. Parameters were extracted from the referenced publication (36).

The extracted parameter values and drug metadata are distributed as three compact SQLite databases. An optional “Ask the data” prototype uses in-browser WebLLM (38) to support natural-language queries against these databases. Entity matching and predefined SQL templates are combined with LLM-assisted query interpretation and constrained SQL generation. Relevant contextual passages are retrieved using BM25 (39) and supplied to the LLM for reference-linked summaries. Results include structured records, provenance, and links to drug-model pages, as illustrated by the metformin clearance query in Figure 9. The query accuracy and usability of this prototype have not been systematically evaluated.

**Figure 9:**
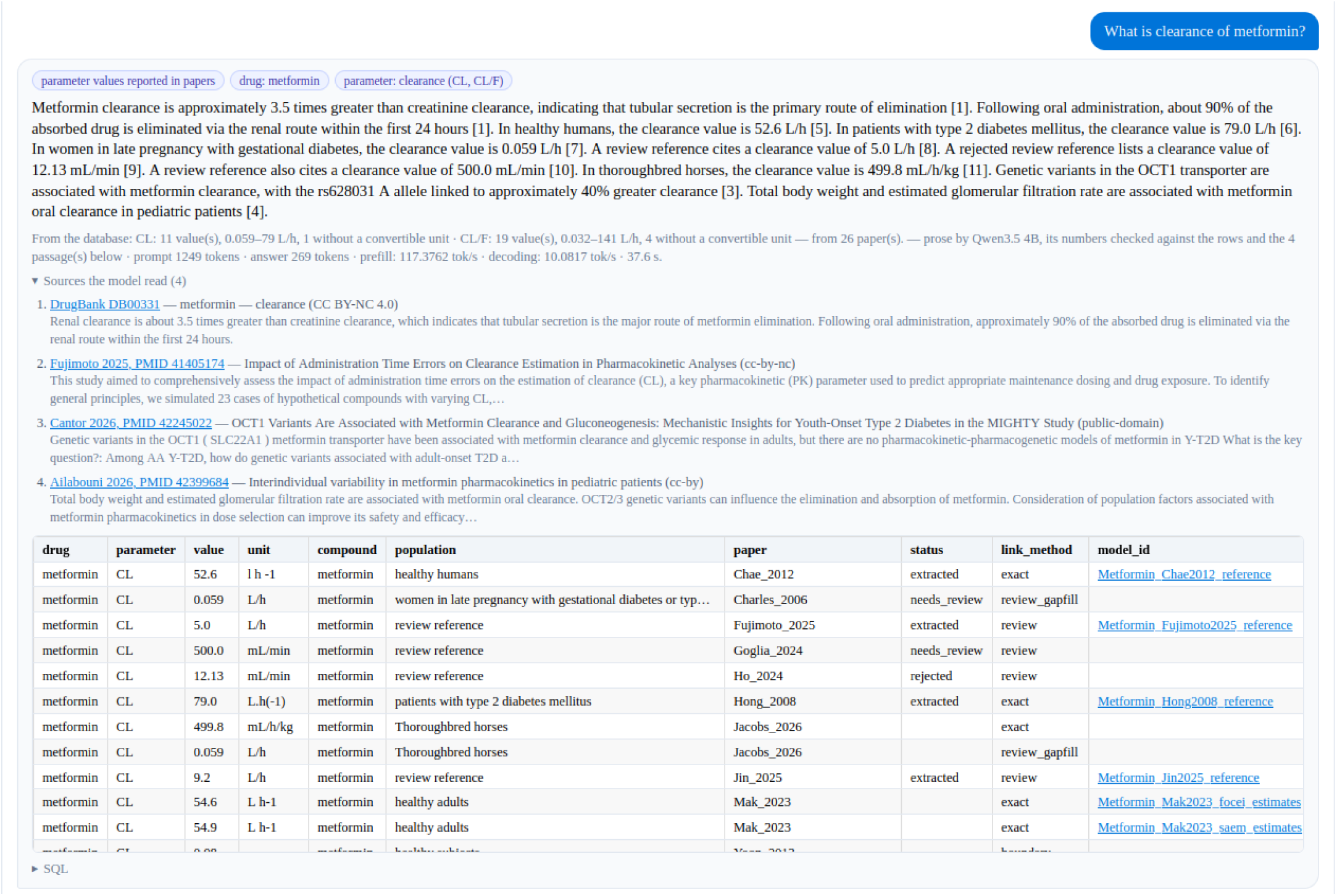
Example of the “Ask the data” prototype responding to a natural-language query about metformin clearance. Retrieved parameter records are presented alongside an LLM-generated summary and contextual information from indexed sources. The table includes values, units, populations, provenance, and review status, with links to the corresponding drug-model pages. This example illustrates the interface functionality rather than a systematic evaluation of response accuracy.

### Knowledge base

The internal knowledge base stores the retrieved literature—including abstracts, full-text PDFs, publisher XML, and supplementary files—together with all intermediate and final artefacts derived from these sources. Its total size currently exceeds ∼200 GB. From this internal store, the public Pharmacolibrary is published as a static website at https://pharmacolibrary.github.io/, with the public Git repository occupying approximately only 0.5 GB.

The public Pharmacolibrary is designed as interlinked database of static web pages so that all computation and querying are performed locally in the browser, using e.g. WebAssembly for simulation and database operations, WebGPU for local LLM inference, and Web Workers for background processing, without requiring server-side computation. The principal practical consideration is therefore the initial download size. Each drug page requires an average download of approximately 121 kB. The models use shared templates, which are translated into shared Functional Mock-up Units (FMUs) of approximately 1 MB each that are cached among multiple models. The special “Ask the data” page downloads additionally approximately 40 MB of compressed SQLite data. If the user selects the small in-browser LLM, approximately 1-4 GB of model weights are downloaded from a third-party provider; these download is optional and remains under the user’s control.

Table 2 summarises statistics collected for a group of 7 selected drugs. This group was processed by the scholar agent’s multi-stage workflow. A PubMed search returned 694 publications, of which 209 were judged relevant after fulltext or abstract-only screening. Of the retrieved publications before judging whether relevant or irrelevant, 216 were obtained as PDF files, either from open-access sources or downloaded manually through an institutional subscription, and converted to a machine-readable structure with GROBID (32); 205 publications were available directly from the publisher in JATS format; 5 were read with the PDFPlumber fallback; and 268 were available as abstracts only. In total, 294 articles were open access and 116 were behind a paywall. In addition, 28 NONMEM control streams and 102 further supplementary files (tables, data, and appendices) were used. The run produced 120 population pharmacokinetics (PK), 108 pharmacodynamic (PD), and 71 pharmacogenomic (PGx) records, from which the engineer agent built 59 executable models. The run took 3 h 40 min, during which the LLM consumed 3.9 million input tokens and generated 535 thousand output tokens.

**Table 2.** One group of drugs processed by the scholar agent’s multi-stage workflow. The workflow ran on a workstation (Intel Core Ultra 7, 8 cores, 16 GB RAM). The LLM (qwen3.8:27b, Q8_0 quantisation, Ollama 0.30.10) ran on a local HPC server (AMD EPYC 7413, 16 dedicated cores, 2 × NVIDIA A100 40 GB).

| drug | time | papers/relevant | full text | access | popPK | PD | PGx | suppl. | review sec. | built | gap-fill | tokens |
|---|---|---|---|---|---|---|---|---|---|---|---|---|
|  |  |  |  |  | ext/rej/other | ext/rej/other | ext/rej/other | NONMEM/other | papers/records | models | pk/pd/pgx | in/out |
| metformin | 1:04:45 | 177/**38** | 27/67/0/83 | 90/4/83 | 23/10/1 | 16/6/0 | 7/0/50 | 6/21 | 40/822 | 15 | **21**/0/– | 396,244/165,135 |
| ibuprofen | 38:35 | 79/**24** | 54/0/2/23 | 13/41/25 | 21/3/1 | 12/25/1 | 8/0/32 | 1/5 | 21/477 | 5 | **13**/**3**/– | 1,016,604/91,654 |
| amoxicillin | 14:43 | 64/**22** | 36/16/2/10 | 23/31/10 | 25/2/2 | 3/4/1 | 0/0/0 | 2/8 | 29/652 | 6 | **34**/0/– | 143,299/33,107 |
| semaglutide | 28:32 | 72/**20** | 5/43/0/24 | 48/0/24 | 13/2/0 | 43/17/2 | 0/0/4 | 7/24 | 19/345 | 7 | **7**/0/– | 144,502/78,047 |
| tolvaptan | 21:08 | 45/**9** | 30/4/0/11 | 17/15/13 | 9/0/0 | 21/3/0 | 2/0/11 | 1/9 | 14/418 | 7 | 0/0/– | 644,865/50,268 |
| allopurinol | 34:48 | 92/**36** | 48/17/1/26 | 40/22/30 | 13/4/1 | 9/12/13 | 40/0/35 | 5/15 | 25/552 | 8 | **15**/0/– | 1,227,333/71,191 |
| clopidogrel | 18:26 | 165/**60** | 16/58/0/91 | 63/3/99 | 16/2/2 | 4/22/0 | 14/0/72 | 6/20 | 37/681 | 11 | **4**/0/– | 123,555/45,513 |
| **TOTAL** | **3:40:59** | **694/209** | **216/205/5/268** | **294/116/284** | 120/23/7 | 108/89/17 | 71/0/204 | **28/102** | **185/3947** | **59** | **94/3/–** | **3,696,402/534,915** |

### Evaluation of extraction accuracy

The benchmark comprised seven drugs selected for therapeutic diversity and the availability of published PGx data: allopurinol, amoxicillin, clopidogrel, ibuprofen, metformin, semaglutide, and tolvaptan. From 24 publications on these drugs, one human expert transcribed a reference set of 146 population PK, PD, and PGx parameter values that the workflow should extract, together with items that it should not extract. This reference set was compared with the output of two variants of the Scholar agent described in Methods:

● Single-pass extraction workflow: with one LLM call per publication
● Multi-stage extraction workflow: with multiple steps with LLM calls and deterministic rules

Both variants use the same publications, the same post-processing that maps parameters to the ontology, and the same scoring; they differ only in the single-pass or multi-stage steps. Benchmark results are in Table 3. The cross-check step was disabled in the benchmark, so each row of Table 3 reflects the output of a single LLM, and the reported token counts and processing times exclude cross-checking.

● **Formatting:** bold rows are benchmark results of multi-stage workflow;
● **TP:** true positives, number of parameters truly present in the referenced publication and extracted with a matching value.
● **FP:** false positives, number of parameters not-present in the referenced publication but wrongly extracted from it.
● **FN:** false negatives, number of parameters present in the referenced publication, but not extracted, or extracted with a wrong value.
● **TN:** true negatives, number of parameters not-present in the referenced publication and correctly not extracted.
● **Metrics:** precision = TP/(TP+FP); recall = TP/(TP+FN); accuracy = (TP+TN)/(TP+FP+FN+TN); F1 = the harmonic mean of precision and recall.

**Table 3.** Extraction performance of different LLMs in the scholar agent (multi-stage vs. single-pass) against the expert reference, sorted by accuracy. Open weighted models wer served by Ollama on a local HPC cluster with 4 available NVIDIA A100 40 GB GPUs; the other rows are OpenAI GPT-5.6 Luna, Terra and Sol, all with high reasoning effort.

| LLM | API | variant | TP | FP | FN | TN | prec | recall | acc | F1 |
|---|---|---|---|---|---|---|---|---|---|---|
| **gpt_5_6_sol__high** | **OpenAI** | **multi-stage** | **92** | **7** | **54** | **168** | **0.929** | **0.63** | **0.81** | **0.751** |
| **qwen3_8_27b_mtp_q8_0** | **Ollama** | **multi-stage** | **87** | **5** | **59** | **168** | **0.946** | **0.596** | **0.799** | **0.731** |
| **gpt_5_6_luna__high** | **OpenAI** | **multi-stage** | **88** | **14** | **58** | **168** | **0.863** | **0.603** | **0.78** | **0.71** |
| **gpt_oss_120b** | **Ollama** | **multi-stage** | **80** | **4** | **66** | **168** | **0.952** | **0.548** | **0.78** | **0.696** |
| **gpt_5_6_terra__high** | **OpenAI** | **multi-stage** | **82** | **15** | **64** | **166** | **0.845** | **0.562** | **0.758** | **0.675** |
| gpt_5_6_sol__high | OpenAI | single-pass | 78 | 16 | 68 | 167 | 0.83 | 0.534 | 0.745 | 0.65 |
| gpt_5_6_luna__high | OpenAI | single-pass | 72 | 11 | 74 | 167 | 0.867 | 0.493 | 0.738 | 0.629 |
| **gemma4_e4b_it_q4_K_M** | **Ollama** | **multi-stage** | **75** | **16** | **71** | **168** | **0.824** | **0.514** | **0.736** | **0.633** |
| gpt_5_6_terra__high | OpenAI | single-pass | 70 | 10 | 76 | 167 | 0.875 | 0.479 | 0.734 | 0.619 |
| gpt_oss_120b | Ollama | single-pass | 69 | 10 | 77 | 168 | 0.873 | 0.473 | 0.731 | 0.613 |
| **qwen3_5_9b** | **Ollama** | **multi-stage** | **86** | **38** | **60** | **174** | **0.694** | **0.589** | **0.726** | **0.637** |
| **gpt_oss_20b** | **Ollama** | **multi-stage** | **64** | **6** | **82** | **168** | **0.914** | **0.438** | **0.725** | **0.593** |
| qwen3_8_27b_mtp_q8 | Ollama | single-pass | 65 | 9 | 81 | 167 | 0.878 | 0.445 | 0.72 | 0.591 |
| **muse_glimmer_30b** | **Ollama** | **multi-stage** | **54** | **7** | **92** | **168** | **0.885** | **0.37** | **0.692** | **0.522** |
| muse_glimmer_30b | Ollama | single-pass | 17 | 11 | 129 | 167 | 0.607 | 0.116 | 0.568 | 0.195 |
| gemma4_e4b_it_q4_K | Ollama | single-pass | 30 | 31 | 116 | 163 | 0.492 | 0.205 | 0.568 | 0.29 |
| qwen3_5_9b | Ollama | single-pass | 1 | 5 | 145 | 168 | 0.167 | 0.007 | 0.53 | 0.013 |

The highest accuracy (81 %) and an F1 score of 0.751 was obtained by the multi-stage variant with OpenAI GPT-5.6 Sol. The open-weights model Qwen3.8:27b, running on the local HPC cluster, reached 80 % with the highest precision among the top models (0.946). In this evaluation, locally deployed open-weights models performed within one percentage point of the best commercial configuration, without sending publications to an external service. With the single-pass variant, the best accuracy was 74 % for GPT-5.6 Sol and 73% for open-weights model gpt-oss:120b.

Across all models for which both variants were evaluated, the multi-stage pipeline outperformed the single-pass workflow, with the largest gains in recall. The largest gains are for small models: Qwen3.5:9b rose from 53 % to 73 %, and the 30B model from 57 % to 69 %. Precision is high throughout (typically above 0.85) while recall stays between 0.37 and 0.63. The dominant error is therefore missed parameters rather than invented ones, suggesting that further development should prioritize parameter-table localization and transcription, while retaining the existing filtering and precision safeguards.

The multi-stage workflow consumed more LLM tokens and took longer than the single-pass workflow: 655 thousand input and 101 thousand output tokens and 23 minutes on average, compared with 250 thousand input and 14 thousand output tokens and 2.2 minutes. It therefore used approximately 2.6 times as many input tokens and seven times as many output tokens, and its wall-clock time was around ten times longer.

## Discussion

Decomposing extraction into smaller subtasks and applying deterministic rules improved performance across all LLMs evaluated with both workflows, enabling open-weight models to achieve results comparable to those of frontier commercial models in this pilot evaluation. Compared with single-pass extraction, accuracy increased by approximately 2–8 percentage points for larger models and by substantially more for some smaller models, at the cost of roughly six– to ninefold longer processing time and threefold higher token usage.

Each parameter is linked to its source, semantic interpretation, model implementation, and simulation result, so the provenance chain can be inspected at every stage of the workflow. Maintaining this chain adds curation effort, particularly when published information is incomplete, ambiguous, or internally inconsistent. The pipeline is therefore not a replacement for expert curation but a means of making large-scale curation systematic, auditable, and incrementally maintainable. The current system supports replication of selected reported quantitative results rather than full reproduction of a published study, which would additionally require its complete data-processing, estimation, and simulation workflow.

Several limitations should be considered when interpreting these results. The benchmark is small, covering seven drugs and 24 publications, and consists mainly of population PK parameters, with few PD and PGx records. The reference set was created by a single human expert and may itself contain interpretation or transcription errors. The pipeline was developed with the Qwen model family, which may favour these models in the comparison. With 146 reference parameters, the differences between the best-performing models are small and should not be interpreted as a ranking. The reference set therefore needs to be extended to more drugs, publications, and record types. Independent curated resources such as nlmixr2lib (5)and the Open Systems Pharmacology model library (9) could serve both as additional reference sets and as sources for verifying model structures and parameters, alongside the NONMEM supplementary materials already used.

Model generation currently covers compartmental pharmacokinetic models with up to three compartments, including selected parent–metabolite structures, and pharmacodynamic models with dose–response or exposure–response behavior. PGx records currently follow the convention of translating activity scores into phenotype classes (poor, intermediate, normal, rapid), which then modify selected covariates of the reference PK model. Future work should extend the workflow to more general structural templates for PK, PD, and PBPK models, and to toxicokinetic and toxicodynamic models, complementing the Open Systems Pharmacology ecosystem (9). Future extensions should also consider genotype effects on PD parameters, transporter-explicit uptake, and continuous activity scores where these are reported.

Pharmacolibrary is not intended as another model repository tied to a single modeling or exchange format; it aims to provide executable models and simulations that do not depend on any one format. Modelica serves as the canonical model representation, and automated cross-implementation tests check that derived implementations preserve the model structure and produce comparable simulation results. Beyond equations, Modelica models can capture complex model structures by connecting reusable components through standardized interfaces, a hierarchical organization that is often lost when models are exported to flat exchange formats. Because these equations and connections are expressed as human-readable plain text, Modelica is also well suited to current LLM-based tools, which can help complete missing components or draft new models from a textual specification; such generated code nevertheless requires the same verification as extracted parameters. Export to other modeling and exchange formats preferred by particular users, together with browser-based simulations, extends the use of source-traceable models beyond a single pharmacometric ecosystem to physiological modeling and education. Interoperability with the nlmixr2 and Open Systems Pharmacology ecosystems, such as importing and exporting models in their native formats, would further make Pharmacolibrary accessible to their established user communities.

The optional “Ask the data” prototype offers natural-language access to structured records and contextual information, but its query accuracy and usability have not been systematically evaluated. Generated summaries should therefore be verified against the linked records and source evidence. Further evaluation is needed to establish the application’s utility for research, education, and clinical information exploration.

Although the knowledge base repository is independently accessible, its generation and scaffolding depend on access to data from external resources whose access policies may change. Continued maintenance therefore requires access to primary or secondary data from life-science literature resources such as Europe PMC (30), PubMed (40), as well as pharmacology-focused resources such as DrugBank (11) or ClinPGx (13,14).

## Conclusion

Pharmacolibrary v26.11 provides a source-traceable collection of executable PK, PD, and PGx models, linking published evidence and extracted parameters through model implementations to simulated behavior, so that discrepancies can be traced to the source data, the model structure, or the implementation. Models are available in several modeling and exchange formats and as browser-based simulations, which reduces dependence on a single software ecosystem. Distribution as a static website under version control keeps infrastructure requirements low and supports incremental, transparent curation.

The pilot evaluation indicates that a multi-stage workflow combining LLM calls with deterministic checks makes automated construction of such a knowledge base feasible, both with commercial LLMs and with medium-sized open-weight models deployed locally. Local deployment is relevant where source materials cannot be sent to external services for confidentiality or data-governance reasons. Human review remains necessary for high-confidence curation; the workflow shifts that effort from repetitive extraction toward ambiguous cases and model discrepancies, with the cross-check by a second LLM indicating where to look.

## Conflict of Interest

None declared.

## Acknowledgements

This work was supported by the European Union’s Horizon Europe projects RAIDO (Grant Agreement No. 101135800) and ONCOSCREEN (No. 101097036) and INQUIRE (Grant Agreement No. 101057499). This work was also supported by the project Digital Twin for Personalized Health (DT4PH) funded by BOSA – Belgian Federal Department of Policy & Support and partially supported by the project VR JIPka, number CZ.01.01.01/01/22_002/0000674 funded by MPO – Ministry of Industry and Trade of the Czech Republic.

Computational resources with open-weights models were provided by the VITO high performance computing cluster and by e-INFRA CZ project (ID:90254), supported by the Ministry of Education, Youth and Sports of the Czech Republic. Shared data storage was provided by CESNET storage facilities supported by the project “e-INFRA CZ” under the programme “Projects of Large Research, Development, and Innovations Infrastructures” LM2023054.

## Notes

https://pharmacolibrary.github.io

https://github.com/pharmacolibrary/pharmacolibrary.github.io

https://github.com/digital-biosystems/Pharmacolibrary

### Competing Interest Statement

The authors have declared no competing interest.

